# Immunization with VLP-based vaccines induces high IgG antibody titers in dermal interstitial fluid

**DOI:** 10.1101/2025.06.25.661557

**Authors:** Alexandra Francian, Srinivasa R. Gadam, Aidan Leyba, Robert M. Taylor, Pavan Muttil, Justin T. Baca, Bryce Chackerian

## Abstract

Although antibodies often exert their effects within the interstitium, the distribution of antibodies within extravascular tissue compartments is not typically measured. Here, IgG titers in peripheral blood and dermal interstitial fluid (ISF) were compared after intramuscular and intradermal immunization with virus-like particle (VLP)-based vaccines. Antibody titers and durability in serum and ISF were similar, indicating that VLP-based vaccines can efficiently elicit strong antibody responses in the skin.

## MAIN TEXT

The potency of vaccines is predominantly determined by measuring antibody levels in circulation. However, antibodies often exert their effects within the interstitial space of a tissue^1,2^. The distribution of antibodies in the extravascular compartments of different tissues is not well established, primarily due to difficulties in obtaining sufficient material to evaluate experimentally^1,3,4^. Interstitial fluid (ISF) exists outside of the vasculature within the extracellular space of tissues. It has a composition similar to blood due to fluid exchange between the vasculature and local interstitial compartments^5^. Several studies cite relatively high protein concentration within ISF, although this appears to be tissue-specific and largely depends on the underlying vasculature^1,3,4^. Ideally, antibody concentration should be measured within the target tissue, but this is not yet feasible for most tissues. Several studies have examined the diversity of immunoglobulins (Ig) in the skin^1,4,6^, including IgM, IgG, IgA, and IgE, but very few have investigated the concentration of antigen-specific IgG following vaccination^7,8^. The skin functions as a barrier to the environment and to potential pathogens. Therefore, it is particularly important to study how the overall protein composition in dermal ISF changes in response to infection, vaccination, or other antigen exposure. This study describes the first longitudinal evaluation of antigen-specific IgG in dermal ISF following immunization with virus-like particle (VLP)-based vaccines.

A variety of microneedle technologies have been developed as promising methods for sampling dermal ISF, and in some cases, may provide alternatives to traditional blood draws using hypodermic needles^1,3,4,6,9,10^. Within the layers of the skin (epidermis, dermis, and hypodermis), the ISF content by volume is highest within the dermis (up to 40%)^5^. Sampling dermal ISF using microneedle arrays is a minimally invasive and less painful alternative to blood sampling through venipuncture. Dermal ISF has been utilized in various clinical applications, including physiological monitoring and diagnostics, particularly for monitoring glucose and other metabolites^10–12^. One advantage of using microneedles over traditional hypodermic needles is that they can be designed to avoid reaching nerve structures or vasculature found in the deeper layers of the dermis and hypodermis, making them minimally invasive^3,4^. Using a simply constructed microneedle apparatus that requires no blistering or suction, we were able to passively collect dermal ISF without causing tissue damage (Fig. 1).

**Figure 1.**
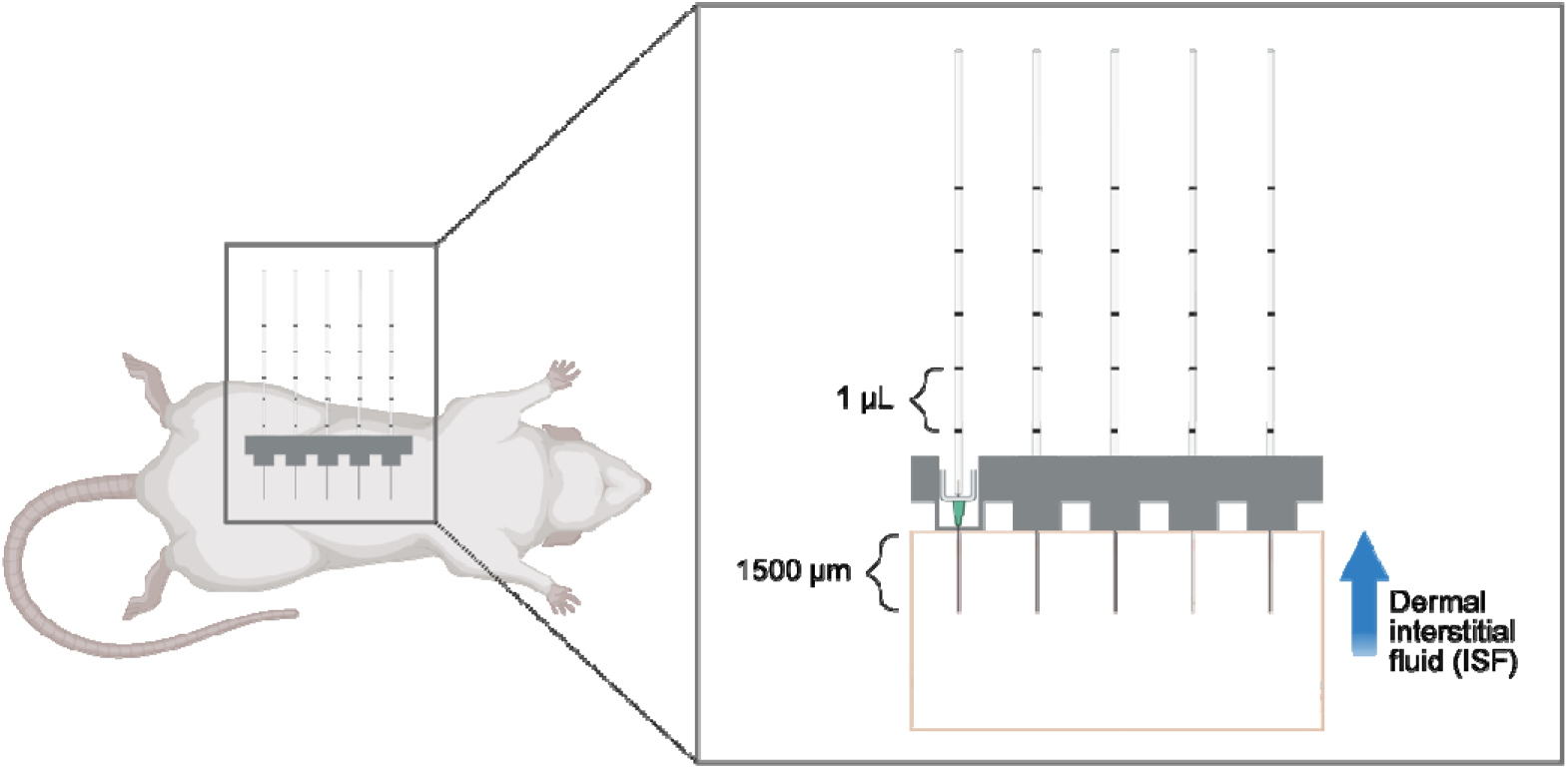
Diagram of microneedle apparatus (MA) for dermal ISF extraction from rats. MAs are 3D printed to hold five ultra-fine nano PEN needles such that 1500 µm of the needle protrudes from the bottom of the array and penetrates the dermis. 5 µL calibrated capillary tubes are placed on the needles inside the MA. MAs are lightly clamped onto the abdomen using 16 mm tissue forceps until at least 10 µL of fluid is collected. ISF samples are visually monitored for blood contamination. Pure ISF samples are transparent to pale yellow; any samples with RBCs present are discarded. Created with BioRender.com.

In this study, we compared antigen-specific IgG titers in dermal ISF with paired serum samples. We evaluated the immunogenicity of three VLP vaccine candidates for mosquito-borne diseases which, ideally, induce antibodies within the dermis (Fig. 2a). These include: 1) L9 VLPs, a malaria vaccine targeting a vulnerable epitope within the circumsporozoite protein (CSP) of *Plasmodium falciparum* sporozoites, the infectious stage of the parasite that causes severe disease in humans^13^. Sporozoites are injected via the bite of *Anopheles* mosquito spp. and traffic through the dermis to the vasculature; 2) TRIO VLPs, a second malaria vaccine targeting the *Anopheles* mosquito spp. salivary protein, TRIO, that was shown to influence mobility and distribution of *P. falciparum* sporozoites within the dermis and exacerbate clinical malaria outcomes^14–16^; and 3) Sialokinin VLPs, a pan-arboviral vaccine targeting a peptide found in *Aedes aegypti* mosquito saliva complicit in enhancing infections caused by multiple mosquito-borne viruses^17,18^. The latter two vaccines are grouped together in this study as “[mosquito] saliva VLPs.” The efficacy of these three vaccines will likely depend on their ability to produce and maintain antibodies at high concentrations in the dermis in order to neutralize their targets quickly; immune memory responses are unlikely to contribute to protection due to the short window of time available for the antibodies to act within the dermis^19,20^. We have previously reported on the engineering, immunogenicity, and efficacy of L9 VLPs^13,21^ and TRIO VLPs^15^. Data describing the sialokinin VLPs are preliminary^22^. Briefly, all three vaccines consist of synthetic peptides conjugated at high valency to the surface of Qβ bacteriophage VLPs (Fig. 2a). Figure 2b demonstrates the method used to calculate conjugation efficiency of peptides to Qβ VLPs, using the sialokinin peptide as an example; each Qβ VLP contains 180 copies of coat protein (coat without peptide attached is 14 kDa, as shown in the WT VLPs in Fig. 2b lane 2). Conjugation efficiency is calculated by a shift in molecular weight upon addition of peptide to coat protein (+1-3 copies of peptide per coat protein, Fig. 2b lane 3). We routinely produce Qβ VLPs displaying more than 300 copies of peptide per particle^13,15^. We have previously shown that all three vaccines induce high-titer and durable antibody responses in mice, with no significant decline in titers throughout their lifespan^13,15,21^ (Fig. 2c, IgG antibody titers from mice immunized with sialokinin VLPs up to 1 year post-immunization). Additionally, L9 VLPs provide significant protection against blood-stage malaria infection in a mouse challenge model, comparable to the WHO-approved RTS,S/AS01_E_ (Mosquirix™; GSK) malaria vaccine^21^. Here, we used a rat model to measure vaccine-elicited antibody titers in dermal ISF and compared these responses with antibody levels in the blood.

**Figure 2.**
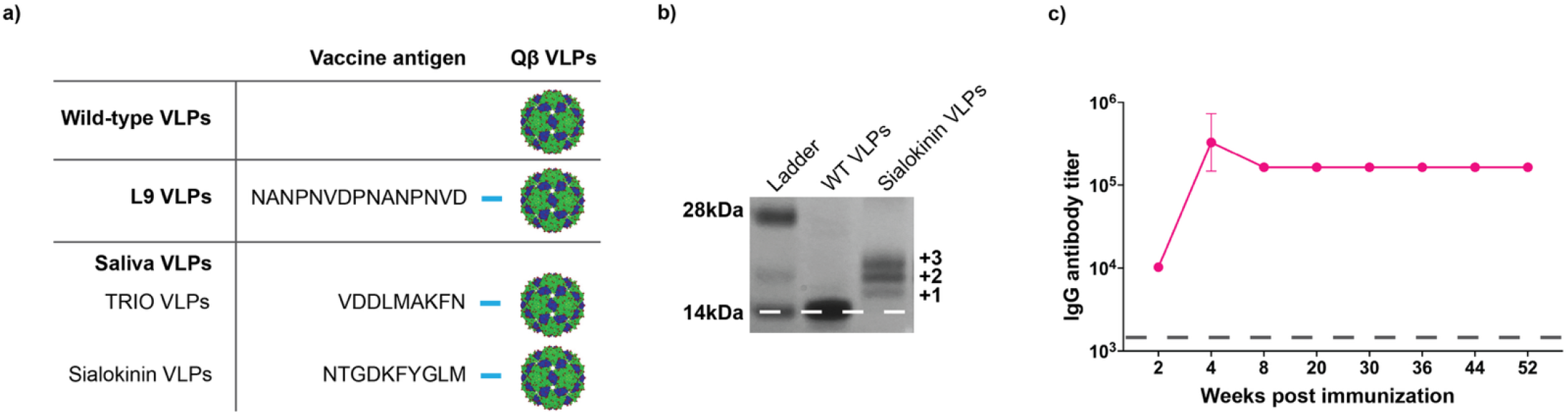
Design of VLP-based vaccines. Three separate VLP-based vaccines are used in this study. **a)** Each vaccine antigen consists of a small peptide synthesized with a four-amino acid linker sequence at the C-terminus (represented by the blue bar). The linker sequence is necessary to chemically conjugate the peptides to purified Qβ VLPs. L9 VLPs display an epitope from the *Plasmodium falciparum* circumsporozoite protein (NANPNVDPNANPNVD), TRIO VLPs display an epitope from the *Anopheles gambiae* salivary protein, TRIO (VDDLMAKFN), and sialokinin VLPs display a ten-amino acid peptide isolated from *Aedes aegypti* saliva (NTGDKFYGLM). **b)** SDS-PAGE analysis of sialokinin-conjugated VLPs. Unmodified Qβ bacteriophage coat protein has a molecular weight of 14 kDa (WT VLPs; lane 2). Conjugation efficiency is assessed via shifts in molecular weight, based on the number of peptides attached per coat protein (+1-3). Gel images are from the same gel. **c)** Durability of IgG antibody response in mice immunized with sialokinin VLPs. Female BALB/c mice (n = 5) were immunized with 5 µg of sialokinin VLPs and boosted 3 weeks later. Blood samples were collected up to 1 year post immunization and IgG antibodies against sialokinin were measured by ELISA. The dashed grey line represents the average IgG titer against WT VLPs (negative control, no antigen) after the second immunization. Data from L9 and TRIO VLP conjugations have been published^13,15,21^

Male and female rats were immunized with 20 µg of L9 VLPs, mosquito saliva VLPs, or WT VLPs (vaccine platform control, no antigen) and boosted 3 weeks later. Rats were immunized intramuscularly or intradermally using dissolvable microneedles^22^. Blood and dermal ISF samples were collected concurrently at 3 timepoints throughout the study: at day 0 prior to immunization, at 5 weeks (2 weeks after the second immunization), and at 6 months (Fig. 3a). Downstream analyses and characterization of vaccine-induced antibody responses in ISF were limited by the small volume of ISF collected (approximately 10 µL per sample). Immunogenicity data in Figure 3 are presented as endpoint dilution titers of IgG antibodies against specific vaccine antigens, measured by ELISA. Figure 3b summarizes the results from rats immunized with L9 VLPs (blue) compared to WT VLPs (black). Titers were measured in sera (circles) and ISF (triangles) against the *Plasmodium falciparum* circumsporozoite protein (CSP) and L9 peptide. Similar to what we previously observed in mice^22^, we saw no difference in antibody responses upon intramuscular or intradermal immunization (Fig. 3c). Therefore, figure 3b shows the combined data from rats that received the same vaccine, regardless of the route of inoculation. L9 VLPs induced significantly higher titers than control WT VLPs in sera and ISF at both timepoints (p values < 0.05). We observed that titers in sera and ISF were mostly indistinguishable from each other. The only significant difference was seen at 5 weeks: rats immunized with L9 VLPs had significantly higher anti-CSP titers in ISF than sera (p value = 0.0286), but this difference was not observed at 6 months. Figure 3d shows IgG titers from rats immunized with TRIO VLPs and sialokinin VLPs (pink) compared to WT VLPs (black). There was no significant difference in titers between the separate mosquito saliva targets, and therefore, the data were combined to provide a larger sample size. We observed similar results to the L9 VLPs: Saliva VLPs induced significantly higher anti-target titers than WT VLPs (p values < 0.05) and there were no significant differences between sera and ISF at both timepoints. In both vaccine groups (Fig. 3b and d), there were no significant decreases in titers between 5 weeks and 6 months (all p values > 0.05), and high IgG titers were sustained in both sera and ISF, indicating that VLP-based vaccines are able to elicit durable antibody reponses in the skin.

**Figure 3:**
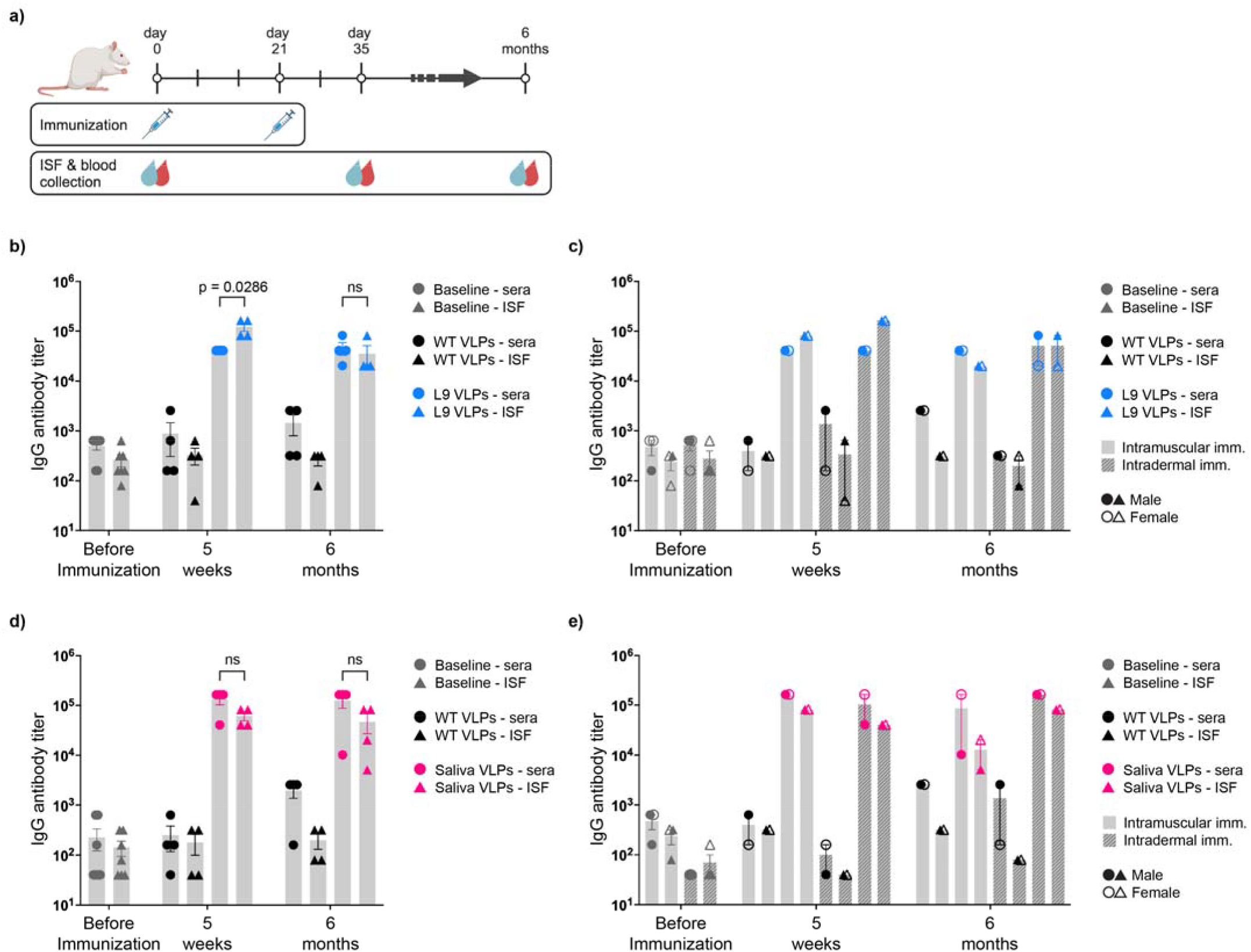
Immunization with VLP-based vaccines results in high IgG antibody titers in sera and interstitial fluid. Male and female Sprague-Dawley rats (n=4; two males and two females per group) were immunized with 20 µg of WT VLPs (negative control, no antigen), L9 VLPs, or mosquito saliva VLPs and boosted 3 weeks later. **a)** Outline of immunizations and sample collection over a 6-month study period. Interstitial fluid (ISF) and blood were collected at day 0 (before immunization; baseline sera and ISF measurements), day 35 (after second immunization), and at 6 months post initial immunization. IgG antibody titers were measured by ELISA against **b, c)** full-length CSP and the L9 peptide from *Plasmodium falciparum*, and **d, e)** two peptides from mosquito saliva, TRIO (*Anopheles gambiae*) and sialokinin (*Aedes aegypti*). **b)** IgG antibody titers against the *Plasmodium falciparum* circumsporozoite protein (CSP) and L9 peptide from rats immunized with L9 VLPs (blue) are compared to titers from rats immunized with WT VLPs (black). Titers were measured in both sera (circles) and ISF (triangles) before immunization, at 5 weeks, and at 6 months. **c)** Data from b) are expanded to show differences between immunization route (solid bar: intramuscular; striped bar: intradermal) and sex (closed shapes: male; open shapes: female). **d)** IgG antibody titers against two mosquito salivary peptides; TRIO from *Anopheles gambiae* mosquitoes and sialokinin from *Aedes aegypti* mosquitoes. Combined titers from rats immunized with saliva VLPs (pink) are compared to titers from rats immunized with WT VLPs (no antigen; black). Titers were measured in both sera (circles) and ISF (triangles) before immunization, at 5 weeks, and at 6 months. **e)** Data from d) are expanded to show differences between immunization route (solid bar: intramuscular; striped bar: intradermal) and sex (closed shapes: male; open shapes: female). **b, d)** All groups had significantly higher titers than WT VLP immunized rats (p values not shown; all p values were < 0.05). Data are reported as mean ± SEM. Significance was determined by one-way ANOVA for multiple comparisons followed by unpaired Mann-Whitney U tests; ns = p value > 0.05. Panel a) was created with BioRender.com.

These results describe the first longitudinal evaluation of antigen-specific IgG in dermal ISF after immunization with VLP-based vaccines. Our analysis revealed equivalent antigen-specific IgG antibodies within sera and ISF for all vaccines tested. Additionally, high IgG titers were sustained in both sera and ISF and did not drop significantly up to 6 months post initial immunization. These results suggest that vaccine-induced IgG concentrations are similar between dermal ISF and serum, adding to results from previous studies that compared protein levels between these compartments ^3,4,6,10,11^ and further supporting the hypothesis that dermal ISF can serve as a proxy for blood in health monitoring applications. These findings have significant translational relevance and could be applied to ISF monitoring in populations with needle-related hesitancies and challenges, pediatric populations, and low resource settings. Future studies should explore the mechanisms of antibody presence and retention in ISF. It was not feasible in this study to measure the exit of antibodies from the skin into the lymphatics or back into circulation, nor did we determine if antibodies detected in ISF simply diffused from circulation or if they are produced locally by skin-resident or skin-associated B cells. Additionally, studies analyzing antibody retention within dermal ISF and the rate at which antibodies traffic between blood, ISF, and lymph would provide a more complete picture on the dynamics of antibody mediated protection within tissue compartments and would be useful for informing antibody dosage and biodistribution studies.

## METHODS

### Qβ virus-like particle (VLP) production

Qβ bacteriophage VLPs are non-infectious particles that are often used as platforms for chemical display. Uniform particles spontaneously assemble when the coat protein is over-expressed in a variety of organisms^23,24^. Coat protein was expressed from the plasmid pETQCT in electrocompetent *Escherichia coli* C41 (DE3) cells (Sigma Aldrich) and isolated from bacterial cell cultures using a standard protein purification protocol^13^, as follows. Bacterial pellets were resuspended in lysis buffer (100 mM NaCl, 10 mM EDTA, 50 mM Tris-HCl, and 0.45% deoxycholate) and incubated on ice for 30 min, followed by three to five cycles of sonication at 20 Hz, until the solution was clear. DNA was removed by adding a solution of 10 µg/mL DNase, 2.5 mM MgCl2, and 0.05 mM CaCl2 (all final concentrations) and incubating on ice for 1 hr. Supernatant was isolated by centrifugation at 10,000 rpm for 30 min (TA-14-50 fixed-angle rotor). Protein was precipitated from supernatant by addition of ammonium sulfate (70% of the final solution) and incubated on ice for 15 min. Precipitated protein was spun at 10,000 rpm for 15 min and the pellet was resuspended in cold SCB buffer (10 mM Tris-HCl, 100 mM NaCl, and 0.1 mM MgSO4). Fractions containing Qβ VLPs were separated by size exclusion chromatography on a Sepharose CL-4B column and identified by gel electrophoresis. Selected fractions were subjected to another round of protein precipitation by 70% ammonium sulfate overnight at 4°C. Precipitated protein was spun at 10,000 rpm for 15 min and pellets were resuspended in phosphate-buffered saline (PBS, pH 7.4). Samples were dialyzed two times against PBS at 4°C. Prior to antigen modification, Qβ VLPs were depleted of bacterial endotoxin by four rounds of phase separation using Triton X-114 (Sigma Aldrich)^25^.

### Conjugation of vaccine antigens to Qβ VLPs

Three separate VLP-based vaccines are used in this study. Each vaccine antigen consists of a small peptide synthesized with a four-amino acid linker sequence at the C-terminus, peptide*-GGGC* (GenScript). The linker sequence is necessary to chemically conjugate the peptides to purified Qβ VLPs. L9 VLPs display the L9 epitope from the *Plasmodium falciparum* circumsporozoite protein (NANPNVDPNANPNVD-*GGGC*), TRIO VLPs display an epitope from the *Anopheles gambiae* salivary protein, TRIO (VDDLMAKFN-*GGGC*), and sialokinin VLPs display a 10 amino acid peptide isolated from *Aedes aegypti* saliva (NTGDKFYGLM-*GGGC*). Peptides are attached to naturally exposed lysine residues on the Qβ VLPs via the heterobifunctional crosslinker, succinyl 6-[(β-maleimidopropionamido)hexanoate] (SMPH; ThermoFisher Scientific). SMPH was incubated with Qβ VLPs at a 10:1 molar ratio (SMPH: Qβ) at room temperature for 2 hrs. Excess SMPH was removed using an Amicon Ultra-4 centrifugal unit with a 100 kDa molecular weight cutoff filter (Millipore Sigma). Peptides were individually added to Qβ VLPs at a 10:1 molar ratio (peptide: Qβ) and incubated overnight at 4°C. Conjugation efficiency was measured via SDS-PAGE, where peptide addition can be visualized by a shift in molecular weight, compared to unconjugated Qβ VLPs. The percentage of coat protein with zero or more attached peptides was determined by SDS-PAGE and used to calculate average peptide density per VLP. Qβ VLPs consist of 180 copies of coat protein per particle, and we consistently produced VLPs displaying more than 300 copies of peptide per particle^13,15^.

### Animals and immunizations

Studies using rats were performed in accordance with the guidelines of the University of New Mexico Animal Care and Use Committee (protocol 22-201289-HSC). A total of twelve 15-20-week-old Sprague-Dawley rats (Crl:SD; Charles River Laboratories, Wilmington, MA, USA) were used for the study. Four rats (two females and two males) were used per group. Rats were immunized intramuscularly or intradermally with 20 µg of VLPs without the addition of exogenous adjuvant. Rats were boosted with the same vaccine 3 weeks after the initial immunization. No differences were seen between immunization routes; therefore, groups receiving the same vaccines were combined to increase sample size from 2 to 4 per group. Rats were immunized with wild-type Qβ VLPs (no antigen, vaccine platform control), L9 VLPs, or mosquito saliva VLPs. Blood and interstitial fluid were collected before immunization, 5 weeks after the initial immunization (2 weeks after the second immunization), and 6 months after the initial immunization.

Animals were anesthetized with 2% Isoflurane (oxygen flow of 1 L/min) before ISF collection. Animals were monitored for signs of distress during ISF extraction and recovery, as well as for one week following the procedure.

### Dermal interstitial fluid (ISF) collection

Dermal ISF was extracted using custom-engineered microneedle arrays (MAs)^10^. MAs were 3D printed to hold five ultra-fine nano PEN needles (Becton Dickinson, NJ, USA). MAs are designed such that 1500 µm of the needle protrudes from the bottom of the array and penetrates the dermis. Several different needle lengths were tested across multiple animals. Briefly, 1000 µm needle length did not collect fluid, 1500 µm reproducibly yielded clear ISF with no blood contamination, and 2000 µm collected blood^10^. Due to MA design and skin deformation during clamping, it is plausible that penetration of the full 1500 µm needle length does not occur; needle length was selected based on ISF collection outcomes and was not confirmed histologically. Calibrated capillary tubes (Drummond Scientific Co., PA, USA) were placed on the needles inside of the MA. MAs were lightly clamped onto the abdomen using 16 mm tissue forceps to provide the necessary pressure for ISF collection (no blistering or suction was required). MAs were applied until at least 10 µL of fluid was collected; if no fluid flow was seen after approximately 10 min, MAs were relocated to a different area on the abdomen. ISF samples were visually monitored for blood contamination. Pure ISF samples are transparent to pale yellow; any samples with RBCs present were discarded. ISF samples were centrifuged to separate any tissue debris and stored at -20°C for subsequent analysis by ELISA.

### Blood collection

Blood samples were collected prior to all ISF collections. Blood was collected by tail snips from anesthetized rats, using non-heparinized capillary blood collection tubes (Kent Scientific Co., CT, USA). Blood was centrifuged to isolate serum, which was then stored at -20°C for subsequent analysis by ELISA.

### Antibody quantification

Antibodies against CSP, L9, TRIO, and sialokinin were detected by ELISA. Recombinant CSP was generously provided by Gabriel Gutierrez (Leidos, Inc.)^26^. Wells of Immulon 2 HB ELISA plates (Thermo Fisher Scientific, Inc.) were coated with 250 ng/well CSP in 50 µL PBS and incubated overnight at 4°C. Wells were blocked with 100 µL blocking buffer (0.5% non-fat dry milk in PBS) per well for 1 h at room temperature. Serum and ISF samples isolated from rats were serially diluted in blocking buffer and applied to wells overnight at 4°C. Reactivity was measured by the addition of HRP-labeled goat anti-rat IgG (Jackson Immunoresearch Laboratories, Inc., PA, USA), diluted 1:5,000 in blocking buffer, and detected by the addition of TMB substrate. Reactions were stopped using 1% HCl and the optical density (OD) of wells was measured at 450 nm.

Peptide ELISAs required an additional step to achieve adequate peptide coating for detection. ELISA plates were coated with 500 ng/well streptavidin at 4°C overnight. SMPH was added at 250 ng/well and incubated for 1 h at room temperature. Wells were then coated with 250 ng/well of peptide (L9, TRIO, or sialokinin) and incubated for 2 h at room temperature. Wells were then blocked, incubated with serial dilutions of sera or ISF, and developed as described above.

### Statistical analyses

Statistical analyses were performed using GraphPad Prism 10 software (version 10.4.2). Statistical significance was determined using one-way ANOVA for multiple comparisons followed by two-tailed, unpaired Mann-Whitney U tests, where a p value < 0.05 was considered significant. Formal power calculations to determine sample size were not possible due to the preliminary nature of this study and modest differences observed may be underpowered due to the small cohort; a retrospective power analysis using G*Power software (version 3.1.9.6) indicated that the minimum sample size to yield a statistical power of at least 0.80 with an alpha of 0.05 and an effect size of 3.66 (calculated from this study) is 3 per group (actual power = 0.88).

## ACKNOWLEDGMENTS

This research was funded by a generous contribution to the UNM Foundation in honor of Jeffrey Michael Gorvetzian in support of biomedical research excellence at the University of New Mexico School of Medicine, by the National Institutes of Health (R01 AI169739 to BC, UL1TR001449 to JTB and RMT, and KL2TR001448 to JTB), and in part by the Dedicated Health Research Funds from the University of New Mexico School of Medicine. AF was supported by the UNM Academic Science Education and Research Training (ASERT) program (funded by K12 GM088021). We also acknowledge facilities provided by the Autophagy, Inflammation, and Metabolism (AIM) Center of Biomedical Research Excellence (COBRE) cores, funded by NIH grant P20 GM121176.

## AUTHOR CONTRIBUTIONS

This work was conceived by AF, BC, RMT, JTB, and PM. AF, SRG, and AL conducted the experiments and AF completed the data analysis. The manuscript was drafted by AF and BC. All authors edited the manuscript and approved its submission.

## COMPETING INTERESTS

BC has equity in Metaphore Biotechnologies. All other authors have no competing interests.

